# Human cytomegalovirus UL2 is not required for lytic replication or viral latency and reactivation

**DOI:** 10.64898/2026.07.30.741858

**Authors:** Amara Bryant, Chris Held, Kamryn Pfenning, Lindsey B. Crawford

## Abstract

Human cytomegalovirus (HCMV) remains a significant cause of morbidity and mortality in transplant patients and is a major cause of congenital disease. Understanding the role of viral genes during infection, viral replication, and viral latency establishment and reactivation is key to the development of new antivirals, which are currently limited. The viral RL11 is a conserved, but understudied viral locus, with previously described roles in specific phases of the viral lifecycle. Antisense to and within the RL11 region are two unrelated genes, including UL2. In this study, we investigate the role of UL2 during viral infection and find that UL2 is not required for lytic replication nor required for viral infection, latency establishment (or persistence) and reactivation in either THP-1 monocytes or primary human hematopoietic progenitor cells (HPCs). As viral genes are evolutionarily optimized for viral fitness, this study suggests that HCMV UL2 has a role outside of viral fitness and may be a unique target or contributor to HCMV-mediated processes including immune response or other viral co-factor regulation.

## Introduction

Human cytomegalovirus (HCMV) is a ubiquitous betaherpesvirus that establishes lifelong persistence following primary infection. In immunocompromised individuals, infection causes significant morbidity and is leading cause of congenital disease (1–6). The viral lifecycle consists of productive lytic infection in permission cells (e.g., fibroblasts, epithelial, endothelial cells) and latent infection of hematopoietic progenitors and myeloid lineage cells, with periodic reactivation enabling continued viral dissemination.

One of the most dynamic regions of the HCMV genome is the RL11 locus. Consisting of the largest and most diverse HCMV gene family, these fourteen genes including UL1 and UL4 through UL11 are predominantly predicted to be glycoproteins and carry an immunoglobulin-like RL11 domain with implications for immune modulation and host adaptation (7, 8). Positioned within this genomic region are two genes, UL2 and UL3, transcribed in antisense orientation to the RL11 locus. Previous large deletion mutants indicated that none of the RL11 proteins, and by default the shared UL2 and UL3 antisense genes, are required for lytic replication in fibroblasts (9–11), although none of these studies evaluated these genes individually. Given the previously demonstrated importance of the RL11 region, including the role of UL4 (12), UL7 (13, 14), and UL8 (15–17) in latency and reactivation of the virus and specific cellular processes controlling viral persistence, we hypothesized that, in combination with, evolutionary conservation of the locus suggests that genes in this region may also contribute to viral fitness. To define the contribution of UL2 to the HCMV lifecycle, we generated a UL2stop mutant virus and assessed viral replication throughout lytic replication in fibroblasts, persistence and reactivation in monocytes, and latency and reactivation in hematopoietic progenitor cells (HPCs).

## Results

### UL2 is conserved across viral strains and encodes a stable viral protein

To establish the level of conservation and expression of UL2, we first compared the coding sequence of UL2 from three commonly used strains of HCMV: TB40/4 BAC4 (EF999921.1), Merlin (NC_006273.2), and AD169 (X17403.1) **(Figure 1A).** Sequence alignment demonstrated 91.2% pair-wise identity between TB40/E and Merlin, and 96.72% pair-wise identity between TB40/E and the lab strain AD169.

**Figure 1.**
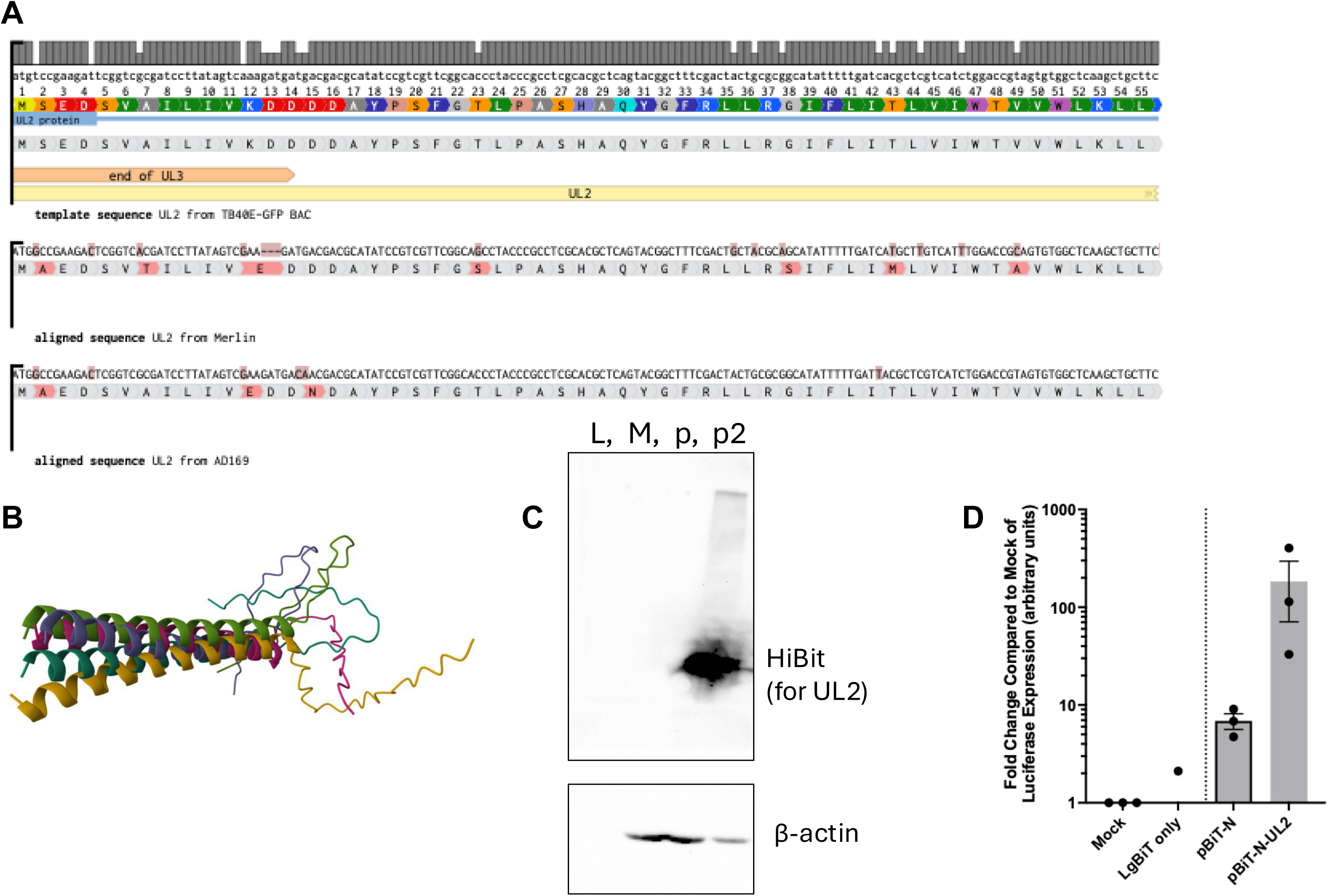
UL2 is a relatively conserved gene and produces an expressed protein. **A)** Sequence comparison of UL2 from three common strains of HCMV: TB40/4 from BAC4, used here (EF999921.1), Merlin (NC_006273.2), and AD169 (X17403.1). **B)** Protein prediction using COSMIC2 version of AlphaFold for six separate runs in overlay. **C)** 293T cells were transfected with a plasmid vector (pBiT-hibit expressing UL2 (p2) using an N-terminal hibit tag) or empty parent plasmid (p) using Lipofectamine 2000 as compared to non-transfected cells (Mock (M)). Cells were harvested in cell lysis buffer (Cell Signaling Technologies) at 48h. Total protein was normalized by Bradford Assay and run on a 7.5% SDS-PAGE gel with sample buffer. Tagged UL2 was detetected with the Promega NanoGlo Hibit Blotting System and total cellular protein using B-actin primary and HRP secondary antibodies. L indicates the molecular weight ladder. D) 293T cells were transfected with Lipofectamine 2000 for pBiT and pBiT-N-UL2 in combination with LgBiT plasmid as in C in white 96-well plates. At 48h, total cells were lysed and HiBiT tagged proteins detected using the Promega NanoGlo HiBiT Lytic Detection System using Luciferase. Data is reported as the relative light units of luciferase for the average of three replicate wells in an experiment normalized to Mock (set to 1) for three independent experiments. Data displayed as mean and SEM for the experimental replicates.

To gain preliminary insight into the structural characteristics of UL2, the predicted protein structure was generated using the TB40/E sequence and the COSMIC2 implementation of AlphaFold. Six prediction runs produced highly similar results (**Figure 1B**), which align with the structure generated by Herpesfolds (18) which used the Merlin reference strain (data not shown). To determine whether UL2 is expressed as a stable protein, we cloned UL2 into the pBiT vector (Promega) which has cloning sites both upstream and downstream of a HiBit tag and uses a CMV IE-based promoter. N-terminal tagged UL2 was transfected into 293T cells in parallel to control empty vector and non-transfected cells. Forty-eight hours post-transfection separate experimental lysates were collected either for analysis of UL2 expression by western blot using a HiBit-specific detection antibody (**Figure 1C**). A specific band at approximately 20kDa was detected only in cells transfected with the UL2 construct, whereas no signal was detected in either Mock- or empty vector-transfected controls. To independently verify UL2 expression, we also analyzed lysates from separate experimental transfections for detection of the HiBit tag using a luciferase-based assay. Total cell lysates were analyzed for luminescence following co-transfection with the LgBit expression plasmid in all groups (**Figure 1D**). Together these data indicate that UL2 encodes a stably expressed viral protein.

### UL2 is dispensable for productive lytic replication in fibroblasts

To determine whether UL2 contributes to productive HCMV infection, we generated a UL2stop mutant in the TB40/E-GFP BAC by introducing three stop codons at positions 5 (Ser->Stop), 18 (Tyr->Stop), and 20 (Ser- >Stop). Due to overlap with the C terminal end of UL3, the first mutation introduces a phenylalanine to leucine mutation at position 98 of UL3, while the remaining mutations are after UL3 termination. Whole BAC sequence analysis demonstrated only the introduced mutations were added (data not shown) and later sequencing of high passage virus post-growth curve analysis demonstrated maintenance of all three mutations (Supplemental Figure 1). HCMV-UL2stop shows no defect in viral reconstitution (data not shown) and no change in viral replication in fibroblasts (**Figure 2**) either in cell-associated (**Figure 2A**) or secreted virus (**Figure 2B**) as measured by infectious titer.

**Figure 2.**
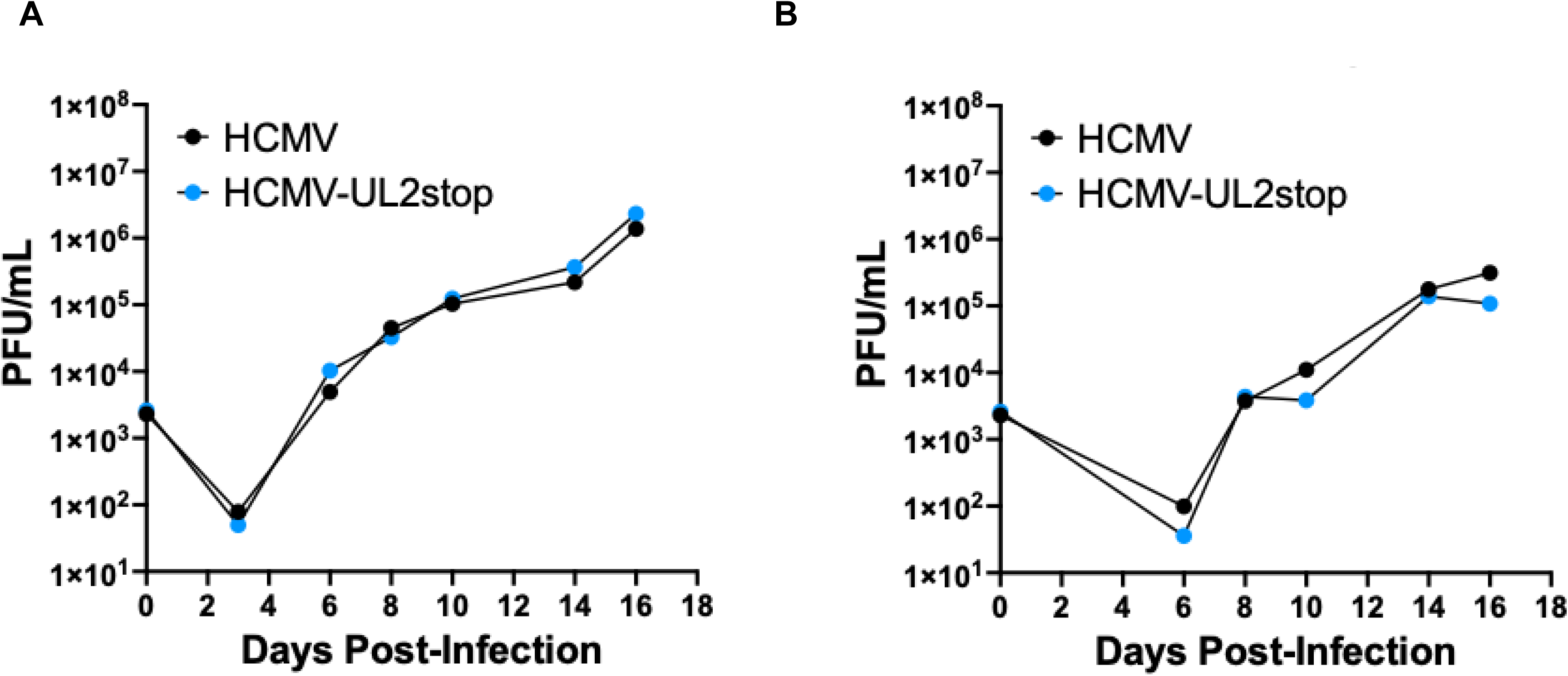
UL2 is not required for replication during lytic infection in fibroblasts. NHDFs were infected with HCMV or HCMV-UL2stop at an MOI of 0.02 for a multi-step growth curve. Cells **(A)** and supernatant **(B)** were collected at the indicated times and stored at −80°C. Viral titer was determined by plaque assay on NHDFs following a 2 week incubation. Data shown is average of two replicate titers per experiment as mean and standard error of the mean for three independent experiments. Individual points for each experiment are shown in Supplemental Figure 2.

### Loss of UL2 does not alter IE1 or UL138 gene expression in THP-1 monocytes

THP-1 monocytes are a common model to study HCMV infection, persistence, and aspects of latency and reactivation. To determine if UL2 plays a role in monocyte infection, persistence, and/or reactivation, THP-1 cells were infected with either wild-type HCMV or HCMV-UL2stop virus and cultured in complete media. Cells were collected at 1, 3, 5, 6, 8, and 11 days post-infection and RNA extracted. To determine viral persistence and monocyte latency, we used qRT-PCR for HCMV IE and UL138, as previously described (19–21). We observed no difference in either lytic (IE, **Figure 3A**) or latent (UL138, **Figure 3B**) markers of infection phase in the absence of UL2 compared to WT virus.

**Figure 3.**
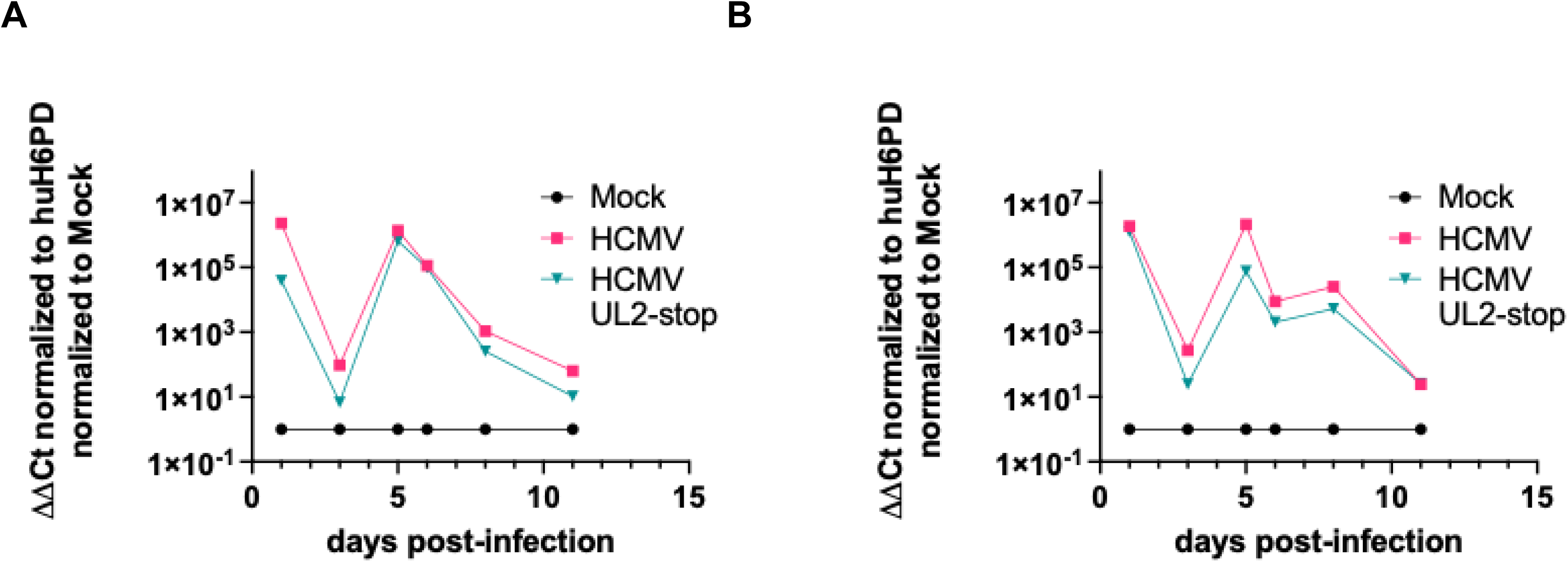
UL2 is not required for latency and reactivation in THP-1 cells. THP-1 monocytes were infected as previously described and cells collected daily. Viral reactivation was induced by TPA treatment at 5dpi with cells and supernatants also collected at 6, 8, 11dpi (=1, 3, 5d post-reactivation). Data shown is representative of 3 independent experiments. RNA was extracted from cells using TRIZOL, cDNA synthesized and qRT-PCR performed for HCMV IE1 **(A)** and HCMV UL138 **(B)** as normalized to human H6PD.

### UL2 is not required for viral reactivation in CD34+ HPCs

Multiple viral genes required for latency and/or reactivation in CD34^+^ hematopoietic progenitor cells (HPCs) are not required for lytic infection or replication and may show specific cell-type specific effects, including those in the surrounding RL11 region (e.g., UL4, UL7, UL8). Therefore, we analyzed whether UL2 is required for latency or reactivation in HPCs. Briefly, as previously described (22), CD34^+^ HPCs were infected with either wild-type HCMV or HCMV-UL2stop virus and cultured for 12 days in a long-term bone marrow culture to maintain CD34^+^ HPC progenitors. Latently infected HPCs were then seeded onto monolayers of permissive fibroblasts in cytokine-rich media to promote myeloid differentiation (12 days latent equivalent to 14 days post-infection). The HPC/fibroblast co-cultures were analyzed weekly by limiting dilution assay for the fraction of GFP^+^ wells for up to four weeks to determine the frequency of infectious centers. As shown in **Figure 4**, loss of UL2 has no effect on the ability of the virus to reactivate. Because we observed no loss of reactivation, we did not further quantify viral genome maintenance in these experiments. These results indicate that UL2 is not required for viral genome maintenance during latency nor for the virus to reactivate in HPCs.

**Figure 4.**
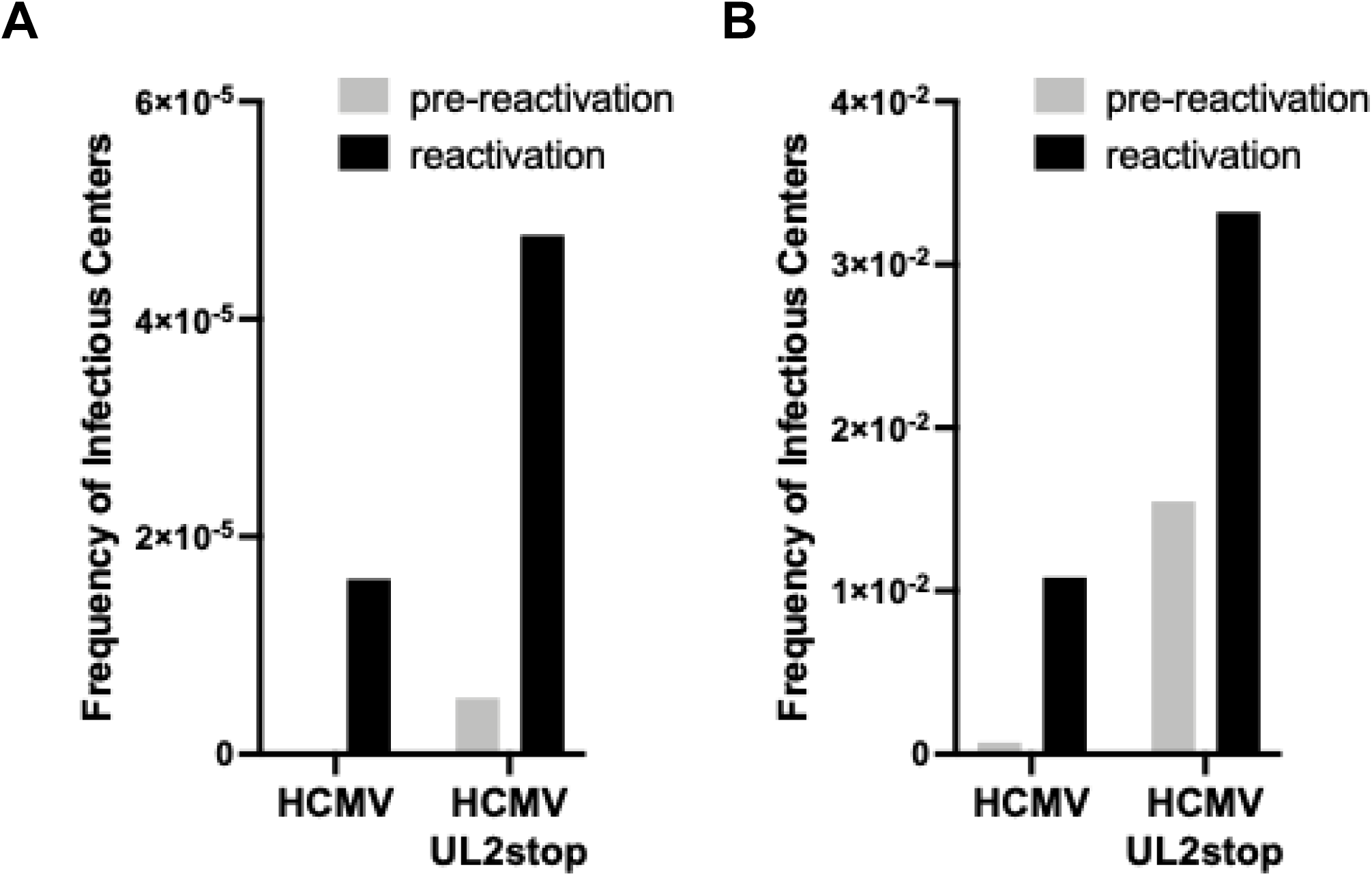
UL2 is not required for latency and reactivation in CD34+ HPCs. **A)** CD34+ HPCs infected with HCMV or HCMV lacking UL2 (HCMV-UL2stop), were sorted, and cultured for latency establishment. At 14dpi (12d latent), equivalent populations of cells were either directly plated on fibroblasts in the presence of cytokine support (reactivation) or lysed and plated on fibroblasts to measure the amount of virus present during latency (pre-reactivation), and the frequency of infectious centers calculated by ELDA. Data shown is the absolute frequency of infectious center formation for 2 independent HPC experiments.

## Discussion

In summary, we identify UL2 as an expressed protein in HCMV that is dispensable for lytic replication in fibroblasts (**Figure 2**) and for IE1 and UL138 expression in THP-1 monocytes (**Figure 3**) and is not required for latency and reactivation in CD34^+^ HPCs (**Figure 4**).

The degree of sequence conservation observed for UL2 between clinical TB40/E and Merlin strains (>90%) and to a highly passaged lab strain AD169 (>97% homology to TB40/E) (**Figure 1**) suggests that UL2 has been maintained despite sequence divergence. While this level of conservation is lower than that observed for core herpesvirus genes (e.g., DNA replication machinery), it is consistent with genes that experience selective pressure and significantly higher than the variable region genes, including some in the immediate neighboring RL11 region (23). Furthermore, expression studies confirm UL2 is a protein coding-region rather than a cryptic or non-functional open reading frame.

Despite protein expression and gene conservation, specific mutation of UL2 to prevent protein expression had no measurable effect on viral replication in fibroblasts (**Figure 2**), suggesting that UL2 is not required for productive lytic infection and suggesting its biological role may be context dependent rather than a core replication component. Several members of the neighboring RL11 gene family contribute specifically to latency and/or reactivation while remaining dispensable for lytic replication (12–17). We therefore investigated whether UL2 functions during latent infection. However, neither latent- nor lytic-associated viral transcripts were altered following infection of a common monocyte model of HCMV latency in THP-1 cells (Figure 3). Nor did loss of UL2 impair viral reactivation from CD34+ hematopoietic progenitor cells, suggesting that UL2 does not play a role in latency establishment or viral genome maintenance during latency, and is dispensable for reactivation. These findings distinguish UL2 from the currently known neighboring RL11 family members with demonstrated roles in latency and/or reactivation.

The apparent lack of a viral maintenance phenotype should not necessarily be interpreted as evidence that UL2 lacks biological function. HCMV encodes numerous accessory proteins whose phenotypes are context dependent, reflecting cell-type specificity (e.g., US28 (24–26) or UL135/UL138), functional redundancy (e.g., US2, US3, US6, US11 (27, 28)), or dependence on coordinated activities of multiple viral proteins or protein isoforms (e.g., UL136 (29–31)). We then hypothesize that UL2 contributes to viral fitness in cellular contexts or stages of infection not examined here including additional hematopoietic or tissue-residence cell populations, modulation of innate or adaptive immune responses, or persistence in vivo.

Transcriptional studies further support the possibility that UL2 has a limited or specifically restricted role during infection. Prior studies detected only low level UL2 transcription in primary hematopoietic cells during latency (32) with little or no expression in clinical latency samples. Additionally, THP-1 latency datasets indicate consistent, but minimal UL2 expression during persistence (19). Additional our transcriptional analysis of fibroblasts infected with WT or UL2stop at 72hpi virus show no significant changes between viruses (data not shown). However, the Weekes lab previously identified 19 cellular proteins that specifically interact with UL2 in a combined overexpression (UL2) and infection (HCMV Merlin strain) proteomics study (33) with a number converging on cellular remodeling and transcriptional and post-transcriptional control of gene expression. This profile suggests that UL2 may function at the interface of cellular architecture and host gene regulation rather than directly targeting classical antiviral signaling pathways.

Together, our findings provide the first functional characterization of HCMV UL2 and demonstrate that UL2 is not required for productive replication, maintenance of latent infection, or reactivation in the experimental systems examined. These findings refine the functional landscape of the greater RL11 gene region by distinguishing UL2 from neighboring genes that play critical roles in latency and reactivation. Future studies examining different cell types, different cellular regulatory functions, and potential redundancy and or negative regulation will be necessary to determine how UL2 contributes to other aspects of HCMV biology that were not assessed here.

## Acknowledgements

This work was supported by Public Health Services grants: P20 GM113126 subproject 6614 and startup funding from the University of Nebraska-Lincoln to LBC. AB was supported in part by a fellowship from the University of Nebraska-Lincoln’s Undergraduate Creative and Research Experience (UCARE) program. We also thank Chloe Deabler for early cloning of UL2 into pCDNA3.1 (not shown). CRediT taxonomy: AB, CH, KP: methodology, investigation, data curation, writing – review & editing; LBC: conceptualization, methodology, investigation, data curation, validation, formal analysis, writing – original draft, writing – review & edition, visualization, supervision, project administration, funding acquisition.

## Materials and Methods

### Cells

Primary CD34^+^ hematopoietic progenitor cells (HPCs) were isolated as previously described (34) or purchased from StemCell Technologies as deidentified samples obtained from human bone marrow. Human embryonic stem cells (NIH approved WA01 and WA09 hESCs) were differentiated into CD34^+^ HPCs as previously described (35). All HPCs were cultured in hematopoietic infection media (IMDM (Hylone) supplemented with 10% BIT 9500 serum substitute (StemCell Technologies), 50 μM 2-mercaptoethanol, 2nM L-glutamine, 20 ng/mL human low-density lipoprotein (Sigma-Aldrich) as previously described (34, 35). THP-1 (bone marrow monoblast), NHDF (normal human dermal fibroblast) cells, and 293T cells were obtained from the American Type Culture Collection (ATCC). THP-1 cells were grown in RPMI-1640 (HyClone or Corning) medium supplemented with 10% fetal bovine serum (FBS) (Gibco), 4.5 g/l glucose, L-glutamine, and antibiotics [penicillin (10 units/ml)–streptomycin (10 ug/ml)]. NHDF and 293T cells were cultured in Dulbecco’s modified Eagle’s medium (DMEM) (Hyclone or Corning) supplemented with 10% FBS, penicillin, streptomycin, and L-glutamine (as above). The M2-10B4 murine stromal cell line expressing human interleukin-3 (IL-3) and granulocyte-colony stimulating factor (G-CSF) and the S1/S1 murine stromal cell line expressing human IL-3 and stem cell factor (SCF) were obtained from Stem Cell Technologies and maintained in DMEM with 10% FBS, penicillin, streptomycin, and L-glutamine as above with selection maintained with G418 and Hygromycin (35).

### Viruses

The HCMV TB40/E bacterial artificial chromosome (BAC) was previously engineered to express green fluorescent protein (GFP) as a visual marker of infection (36). Recombinant HCMV TB40/E-UL2stop was constructed by two-step GalK-Kan recombination (37) using pC255 as a template to generate the GalK-Kan insert and the primers HCMV-UL2-GK-F forward (5’-CTCACTATCCCCCGAGCCGGAGCCGAACGATTCCAAACCGGATACGCTACATACCAGCCACAGTGGGTGGCTTTTCCTGTTGACAATTAATCATCGGCA-3’) and HCMV-UL2-GK-R reverse (5’-TGAAGGAAAGTACAGGTTCCTGTTGATATGTTATTACAGAAGGTCACGGAACACAAACGTTTTCTGCGTGTGTCTCAGCAAAAGTTCGATTTATTCAAC-3’). Following selection of recombinant BACs on the basis of kanamycin resistance, the Kan^R^ marker was removed using minimal DOG selection. Recombination was assessed at each step using flanking primers: HCMV-UL2-flanking-F forward (5’-CACCATGTGCCTCATCTGTC-3’) and HCMV-UL3-flanking-R reverse (5’-TTTGCAAATCAGCAGAGACG-3’) at each step for the removal of wild-type sequence and insertion of GK-Kan insert and removal of the GK-Kan insert and insertion of the UL2stop region, respectively. UL2stop was generated with recombination primers: HCMV-UL2stop-F forward (5’-ATACGCTACATACCAGCCACAGTGGGTGGCTTTTACCATGTCCGAAGATTAGGTCGCGATCCTTATAGTCAAAGATGATGACGACGCATAACCGTAGTTC-3’) and HCMV-UL2stop-R reverse (5’- TGAAGGAAAGTACAGGTTCCTGTTGATATGTTATTACAGAAGGTCACGGAACACAAACGTTTTCTGCGTGTGTTTTTATAAAAGAGCGTCTCGAAGCAGC-3’). BAC integrity was examined by enzyme digest analysis and sequencing of the entire viral genome by next generation sequencing. Virus was reconstituted by transfecting the BAC genomes (1 µg) into NHDF fibroblasts and incubating until >80% cytopathic effects (CPE) were observed. Plaque purification was performed and HCMV TB40/E-GFP-UL2stop stocks and titers were generated as previously described (38). UL2stop mutation was assessed by 1) whole BAC sequencing at time of reconstitution and 2) periodic viral genome sequencing to confirm retention of the mutation on later viral stock passages or following growth curve analysis was performed by PCR product sequencing. A PCR product of the region from outside UL2 through outside UL9 of ∼5kB (using primers: HCMV_UL2_flanking_F: 5’-CACCATGTGCCTCATCTGTC-3’, and HCMV_UL9_flanking_R: 5’-CATGGAAAAAGGCCATGACT-3’) was amplified using the Expand Long-template PCR kit (Roche) and sequenced by PlasmidSaurus. All viral mutants were confirmed to be maintained through at least passage 7 (virus used between passages 4 and 6 for all experiments). Viruses were titered in parallel (UL2stop and WT) by plaque assay for all experiments.

### Immunoblotting

Extracts were run on 7.5% SDS-PAGE, transferred to PDVF membrane and visualized with antibodies specific for HiBit tag using the NanoGlo HiBiT blotting system (Promega) following the manufacturer’s instructions. Cellular controls were stained on stripped blots probed with B-actin (Cell Signaling Technologies cl.D6A8) followed by Rabbit anti-Mouse HRP (Cell Signaling Technologies).

### Latency and Reactivation

CD34^+^ HPCs were infected at an MOI of 3 for 42 h. Following infection, pure populations of viable, infected (GFP-positive) CD34^+^ HPCs were isolated by fluorescence-activated cell sorting (FACS) using an Aria II (Becton Dickson) using PI for viability and an APC-conjugated CD34 antibody (Biolegend). Long-term cultures were maintained in Myelocult supplemented with hydrocortisone in transwells above an irradiated M2-10B4 and S1/S1 stromal cell monolayer for 12 days. The frequency of infectious center production during the culture period was measured using a limiting dilution assay (39). HCMV latency and reactivation protocols are detailed in depth in (35).

### Quantitative (q) PCR for Viral Genomes and qRT-PCR for Viral Gene Expression

Total DNA and RNA were extracted from cells using TRIZOL manufacturer’s directions. Primers and a probe recognizing HCMV UL141 were used to quantify HCMV genomes (probe = CGAGGGAGAGCAAGTT; forward primer = 5’ GATGTGGGCCGAGAATTATGA and reverse primer = 5’ ATGGGCCAGGAGTGTGTCA) as previously described. Viral genomes synthesized during infection in CD34^+^ HPCs were normalized to total cell number determined using human β-globin as a reference (probe = GGACAGATCCCCAAAGGACT; forward primer = 5’ TTAGGGTTGCCCATAACAGC and reverse primer = 5’ TTGGACCCAGAGGTTCTTTG) as previously described using TaqMan FastAdvanced. IE1 and UL138 were detected using SybrGreen qRT-PCR (Abcam) and previously published primers (40).

### Statistical Analysis

Statistical analysis was performed using GraphPad Prism (v9) for comparison between groups using an unpaired Student’s t-test or one-way Anova or two-way Anova with p values as indicated in each figure.

## Figure Legends

**Supplemental Figure 1.**
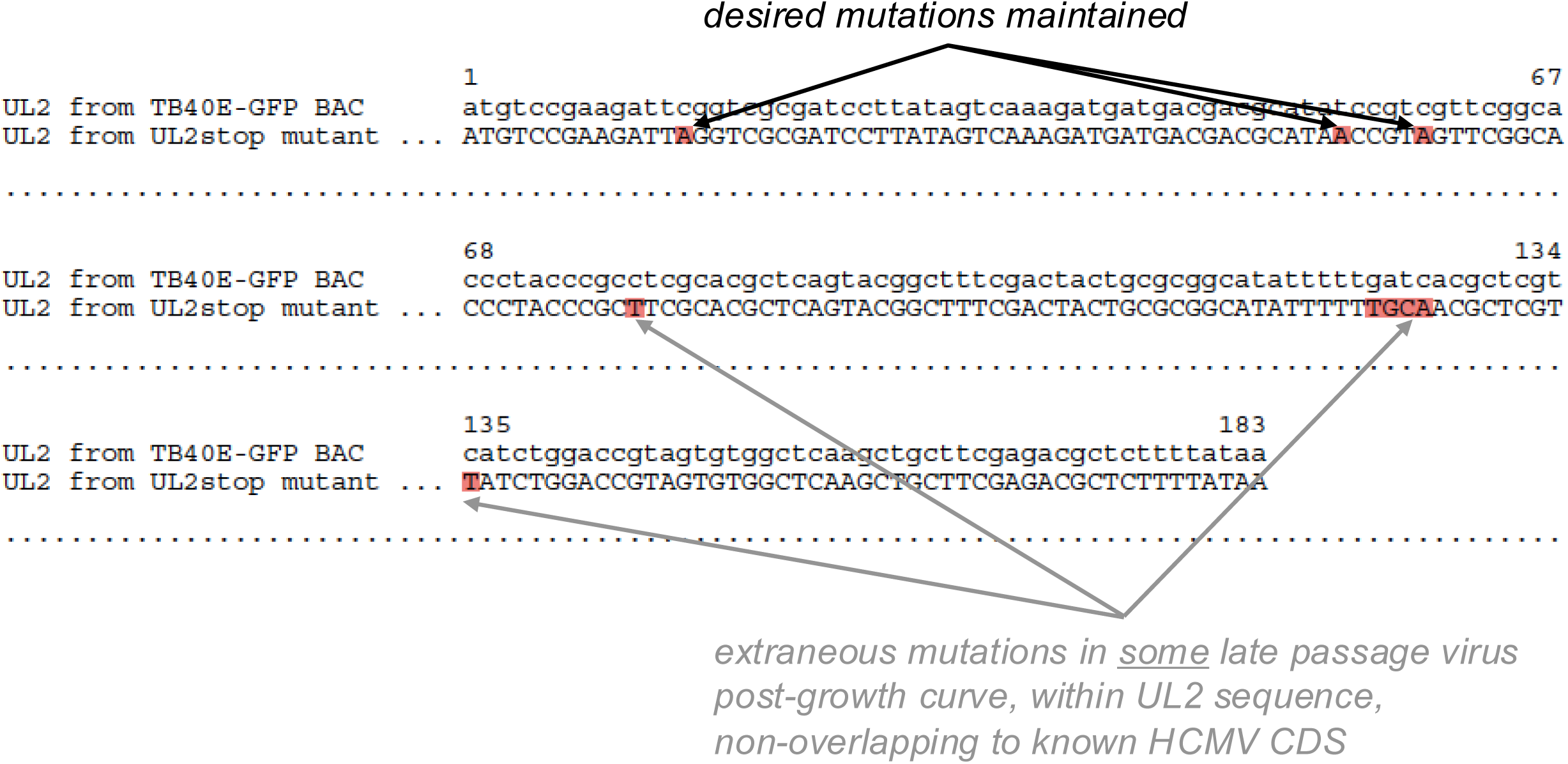
Sequence Analysis of the UL2 region post-growth curve in late passage virus. Sequence comparison of the UL2 coding region as amplified from viral DNA (from supernatant from day 16 of a low MOI growth curve for the HCMV-UL2stop mutant virus), amplified by PCR for the UL2 through UL9 region and sequenced as compared to the TB40/E reference sequence. WT viral DNA from the same experiment and the remaining non-UL2 region for UL2 through UL9 were identical to the reference sequence.

**Supplemental Figure 2.**
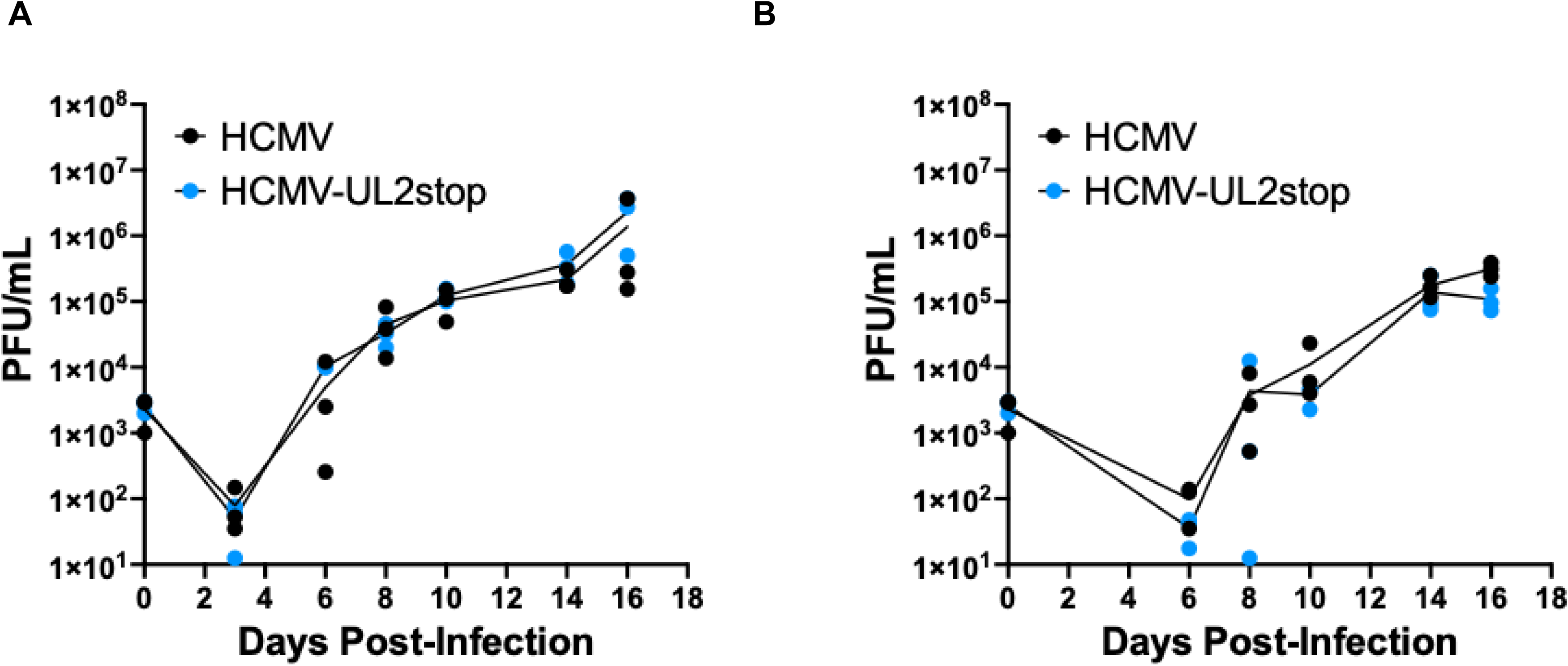
Individual Experimental Growth Curve Data from Figure 2. NHDFs were infected with HCMV or HCMV-UL2stop at an MOI of 0.02 for a multi-step growth curve. Cells **(A)** and supernatant **(B)** were collected at the indicated times and stored at −80°C. Viral titer was determined by plaque assay on NHDFs following a 2 week incubation. Data shown is average of two replicate titers per experiment with each experiment visible and no error bars for all three independent experiments. Data with statistical analysis (mean with SEM for experimental replicates) are shown in Figure 2.

## References

1. Ramanan P, Razonable RR. 2013. Cytomegalovirus infections in solid organ transplantation: a review. Infect Chemother 45:260–71.

2. Nogalski MT, Collins-McMillen D, Yurochko AD. 2014. Overview of human cytomegalovirus pathogenesis. Methods Mol Biol 1119:15–28.

3. Voigt S, Schaffrath Rosario A, Mankertz A. 2016. Cytomegalovirus Seroprevalence Among Children and Adolescents in Germany: Data From the German Health Interview and Examination Survey for Children and Adolescents (KiGGS), 2003-2006. Open Forum Infect Dis 3:ofv193.

4. Stern L, Withers B, Avdic S, Gottlieb D, Abendroth A, Blyth E, Slobedman B. 2019. Human Cytomegalovirus Latency and Reactivation in Allogeneic Hematopoietic Stem Cell Transplant Recipients. Front Microbiol 10:1186.

5. Xia W, Yan H, Zhang Y, Wang C, Gao W, Lv C, Wang W, Liu Z. 2021. Congenital Human Cytomegalovirus Infection Inducing Sensorineural Hearing Loss. Frontiers in microbiology 12:649690–649690.

6. Annaloro C, Serpenti F, Saporiti G, Galassi G, Cavallaro F, Grifoni F, Goldaniga M, Baldini L, Onida F. 2021. Viral Infections in HSCT: Detection, Monitoring, Clinical Management, and Immunologic Implications. Frontiers in immunology 11:569381–569381.

7. Davison AJ, Akter P, Cunningham C, Dolan A, Addison C, Dargan DJ, Hassan-Walker AF, Emery VC, Griffiths PD, Wilkinson GWG. 2003. Homology between the human cytomegalovirus RL11 gene family and human adenovirus E3 genes. J Gen Virol 84:657–663.

8. Litvin U, Wang ECY, Stanton RJ, Fielding CA, Hughes J. 2024. Evolution of the Cytomegalovirus RL11 gene family in Old World monkeys and Great Apes. Virus Evol 10:veae066.

9. Hobom U, Brune W, Messerle M, Hahn G, Koszinowski UH. 2000. Fast screening procedures for random transposon libraries of cloned herpesvirus genomes: mutational analysis of human cytomegalovirus envelope glycoprotein genes. Journal of virology 74:7720–7729.

10. Yu D, Smith GA, Enquist LW, Shenk T. 2002. Construction of a self-excisable bacterial artificial chromosome containing the human cytomegalovirus genome and mutagenesis of the diploid TRL/IRL13 gene. Journal of virology 76:2316–2328.

11. Atalay R, Zimmermann A, Wagner M, Borst E, Benz C, Messerle M, Hengel H. 2002. Identification and expression of human cytomegalovirus transcription units coding for two distinct Fcgamma receptor homologs. Journal of virology 76:8596–8608.

12. Held C, Pfenning K, Crawford LB. 2026. Human Cytomegalovirus UL4 is Required for Viral Reactivation via Cellular Reprogramming in Latently Infected CD34+ Progenitor Cells and Humanized Mice. bioRxiv doi:10.64898/2026.06.28.735103:2026.06.28.735103.

13. Crawford LB, Kim JH, Collins-McMillen D, Lee BJ, Landais I, Held C, Nelson JA, Yurochko AD, Caposio P. 2018. Human Cytomegalovirus Encodes a Novel FLT3 Receptor Ligand Necessary for Hematopoietic Cell Differentiation and Viral Reactivation. mBio 9.

14. Hancock MH, Crawford LB, Perez W, Struthers HM, Mitchell J, Caposio P. 2021. Human Cytomegalovirus UL7, miR-US5-1, and miR-UL112-3p Inactivation of FOXO3a Protects CD34(+) Hematopoietic Progenitor Cells from Apoptosis. mSphere 6.

15. Dirck A, Diggins NL, Crawford LB, Perez WD, Parkins CJ, Struthers HH, Turner R, Pham AH, Mitchell J, Papen CR, Malouli D, Hancock MH, Caposio P. 2023. HCMV UL8 interaction with β-catenin and DVL2 regulates viral reactivation in CD34(+) hematopoietic progenitor cells. J Virol 97:e0124123.

16. Dirck A, Diggins N, Perez W, Parkins C, Daily M, Turner R, Slind L, Nguyen L, Malouli D, Wu G, Hancock M, Caposio P. 2025. HCMV promotes viral reactivation through the coordinated regulation of Notch signaling by UL8 and miR-UL36. bioRxiv doi:10.1101/2025.11.05.686718.

17. Dirck A, Diggins N, Perez W, Parkins C, Daily M, Turner RL, Slind L, Nguyen L, Malouli D, Wu G, Hancock M, Caposio P. 2026. HCMV promotes viral reactivation through the coordinated regulation of Notch signaling by UL8 and miR-UL36. mBio doi:10.1128/mbio.03377-25:e0337725.

18. Soh TK, Ognibene S, Sanders S, Schäper R, Kaufer BB, Bosse JB. 2024. A proteome-wide structural systems approach reveals insights into protein families of all human herpesviruses. Nature Communications 15:10230.

19. Collins-McMillen D, De Oliveira Pessoa D, Zarrella K, Parkins CJ, Daily M, McKinzey DR, Moorman NJ, Kamil JP, Caposio P, Padi M, Goodrum FD. 2025. Viral and host network analysis of the human cytomegalovirus transcriptome in latency. Proc Natl Acad Sci U S A 122:e2416114122.

20. Krishna BA, Poole EL, Jackson SE, Smit MJ, Wills MR, Sinclair JH. 2017. Latency-Associated Expression of Human Cytomegalovirus US28 Attenuates Cell Signaling Pathways To Maintain Latent Infection. mBio 8:e01754–17.

21. Lau B, Poole E, Van Damme E, Bunkens L, Sowash M, King H, Murphy E, Wills M, Van Loock M, Sinclair J. 2016. Human cytomegalovirus miR-UL112-1 promotes the down-regulation of viral immediate early-gene expression during latency to prevent T-cell recognition of latently infected cells. J Gen Virol 97:2387–2398.

22. Umashankar M, Goodrum F. 2014. Hematopoietic Long-Term Culture (hLTC) for Human Cytomegalovirus Latency and Reactivation, p 99-112. *In* Yurochko AD, Miller WE (ed), Human Cytomegaloviruses, vol 1119. Humana Press.

23. Lehner R, Stamminger T, Mach M. 1991. Comparative sequence analysis of human cytomegalovirus strains. J Clin Microbiol 29:2494–502.

24. Krishna BA, Wass AB, Sridharan R, O’Connor CM. 2020. The Requirement for US28 During Cytomegalovirus Latency Is Independent of US27 and US29 Gene Expression. Front Cell Infect Microbiol 10:186.

25. Krishna BA, Humby MS, Miller WE, O’Connor CM. 2019. Human cytomegalovirus G protein-coupled receptor US28 promotes latency by attenuating c-fos. Proc Natl Acad Sci U S A 116:1755–1764.

26. Crawford LB, Caposio P, Kreklywich C, Pham AH, Hancock MH, Jones TA, Smith PP, Yurochko AD, Nelson JA, Streblow DN. 2019. Human Cytomegalovirus US28 Ligand Binding Activity Is Required for Latency in CD34(+) Hematopoietic Progenitor Cells and Humanized NSG Mice. mBio 10.

27. Rehm A, Engelsberg A, Tortorella D, Körner IJ, Lehmann I, Ploegh HL, Höpken UE. 2002. Human cytomegalovirus gene products US2 and US11 differ in their ability to attack major histocompatibility class I heavy chains in dendritic cells. J Virol 76:5043–50.

28. Pande NT, Powers C, Ahn K, Früh K. 2005. Rhesus cytomegalovirus contains functional homologues of US2, US3, US6, and US11. J Virol 79:5786–98.

29. Moy MA, Collins-McMillen D, Crawford L, Parkins C, Zeltzer S, Caviness K, Zaidi SSA, Caposio P, Goodrum F. 2023. Stabilization of the human cytomegalovirus UL136p33 reactivation determinant overcomes the requirement for UL135 for replication in hematopoietic cells. J Virol 97:e0014823.

30. Moy MA, Collins-McMillen D, Crawford L, Parkins C, Zeltzer S, Caviness K, Caposio P, Goodrum F. 2023. UL135 and UL136 Epistasis Controls Reactivation of Human Cytomegalovirus. bioRxiv doi:10.1101/2023.01.24.525282:2023.01.24.525282.

31. Caviness K, Bughio F, Crawford LB, Streblow DN, Nelson JA, Caposio P, Goodrum F. 2016. Complex Interplay of the UL136 Isoforms Balances Cytomegalovirus Replication and Latency. mBio 7:e01986.

32. Cheng S, Caviness K, Buehler J, Smithey M, Nikolich-Zugich J, Goodrum F. 2017. Transcriptome-wide characterization of human cytomegalovirus in natural infection and experimental latency. Proc Natl Acad Sci U S A 114:E10586–E10595.

33. Nobre LV, Nightingale K, Ravenhill BJ, Antrobus R, Soday L, Nichols J, Davies JA, Seirafian S, Wang EC, Davison AJ, Wilkinson GW, Stanton RJ, Huttlin EL, Weekes MP. 2019. Human cytomegalovirus interactome analysis identifies degradation hubs, domain associations and viral protein functions. Elife 8:e49894.

34. Crawford LB, Caposio P. 2021. Development of a huBLT Mouse Model to Study HCMV Latency, Reactivation, and Immune Response. Methods Mol Biol 2244:343–363.

35. Crawford LB. 2022. Human Embryonic Stem Cells as a Model for Hematopoietic Stem Cell Differentiation and Viral Infection. Curr Protoc 2:e622.

36. Umashankar M, Petrucelli A, Cicchini L, Caposio P, Kreklywich CN, Rak M, Bughio F, Goldman DC, Hamlin KL, Nelson JA, Fleming WH, Streblow DN, Goodrum F. 2011. A novel human cytomegalovirus locus modulates cell type-specific outcomes of infection. PLoS Pathog 7:e1002444.

37. Paredes AM, Yu D. 2012. Human cytomegalovirus: bacterial artificial chromosome (BAC) cloning and genetic manipulation. Curr Protoc Microbiol Chapter 14:Unit14E.4.

38. MacManiman JD, Meuser A, Botto S, Smith PP, Liu F, Jarvis MA, Nelson JA, Caposio P. 2014. Human Cytomegalovirus-Encoded pUL7 Is a Novel CEACAM1-Like Molecule Responsible for Promotion of Angiogenesis. mBio 5.

39. Hu Y, Smyth GK. 2009. ELDA: extreme limiting dilution analysis for comparing depleted and enriched populations in stem cell and other assays. J Immunol Methods 347:70–8.

40. Zarrella K, Longmire P, Zeltzer S, Collins-McMillen D, Hancock M, Buehler J, Reitsma JM, Terhune SS, Nelson JA, Goodrum F. 2023. Human cytomegalovirus UL138 interaction with USP1 activates STAT1 in infection. PLoS Pathog 19:e1011185.

